# Loss of SATB1 Induces a p21 Dependent Cellular Senescence Phenotype in Dopaminergic Neurons

**DOI:** 10.1101/452243

**Authors:** Markus Riessland, Benjamin Kolisnyk, Tae Wan Kim, Jia Cheng, Jason Ni, Jordan A. Pearson, Emily J. Park, Kevin Dam, Devrim Acehan, Lavoisier S. Ramos-Espiritu, Wei Wang, Jack Zhang, Jae-won Shim, Gabriele Ciceri, Lars Brichta, Lorenz Studer, Paul Greengard

**Author notes:** contributed equally (Markus Riessland, Benjamin Kolisnyk, Tae Wan Kim). These authors contributed equally to directing this work. Correspondence should be addressed to P.G. or L.S.

## Abstract

Cellular senescence is a mechanism used by mitotic cells to prevent uncontrolled cell division. As senescent cells persist in tissues, they cause local inflammation and are harmful to surrounding cells, contributing to aging. Generally, neurodegenerative diseases, such as Parkinson‘s, are disorders of aging. The contribution of cellular senescence to neurodegeneration is still unclear. SATB1 is a DNA binding protein associated with Parkinson’s disease. We report that SATB1 prevents cellular senescence in post-mitotic dopaminergic neurons. Loss of SATB1 causes activation of a cellular senescence transcriptional program in dopamine neurons, both in human stem cell-derived dopaminergic neurons and in mice. We observed phenotypes which are central to cellular senescence in SATB1 knockout dopamine neurons *in vitro* and *in vivo*. Moreover, we found that SATB1 directly represses expression of the pro-senescence factor, p21, in dopaminergic neurons. Our data implicate senescence of dopamine neurons as a contributing factor to the pathology of Parkinson’s disease.

## Introduction

A key pathological hallmark of Parkinson‘s disease (PD) is the progressive degeneration of dopaminergic (DA) neurons in the substantia nigra pars compacta (SNpc) of the midbrain. The loss of these cells causes the characteristic motor symptoms of the disease. In contrast, non-dopaminergic neuronal populations, such as cortical (CTX) neurons, remain generally spared in the disease (Giguere et al., 2018; Kalia and Lang, 2015; Pedersen et al., 2005; Rajput and Rozdilsky, 1976). The molecular basis of PD remains elusive. Identification of causative Parkinson’s genes has shed light on the molecular mechanisms of the degeneration of DA neurons. The functions of many PD genes converge upon mitochondrial quality control (PINK1, PRKN) and lysosomal function (GBA) (Mullin and Schapira, 2015). Age, however, remains the principal risk factor in both the sporadic and familial forms of the disease (Reeve et al., 2014). In line with this, mitochondrial and lysosomal dysfunction are both linked with aging (Folick et al., 2015; Tauchi and Sato, 1968).

Interestingly, aging organisms accumulate senescent cells within their tissues (Jeyapalan et al., 2007). Although detrimental later in life, senescence is an important process in both embryogenesis and wound healing and thus is beneficial early in life (Storer and Keyes, 2014). Senescent cells have lysosomal and mitochondrial abnormalities. They can damage or negatively impact surrounding cells through the senescence associated secretory phenotype (SASP) (Acosta et al., 2013). Elimination of senescent cells increases both healthspan and lifespan (Xu et al., 2018). Besides healthy aged tissue, senescent cells also accumulate in many disease states. It has recently been reported that there is an increase in cellular senescence markers in the midbrain of PD patients (Chinta et al., 2018). Given that cellular senescence is characteristic of mitotic cells, the upregulation of senescence markers in the CNS was therefore attributed to the mitotic cells of the brain. There is however growing interest in cellular senescence in post-mitotic cells (Sapieha and Mallette, 2018). Multiple lines of evidence suggest that terminally differentiated neurons may shift to a state similar to senescence (Baker and Petersen, 2018; Jurk et al., 2012; Piechota et al., 2016).

Cellular senescence is a mechanism of cell cycle arrest, to protect from uncontrolled proliferation (Campisi, 2013; Kirkland and Tchkonia, 2017). Both DNA damage and oxidative stress are able to induce cellular senescence. Two pathways both initiate and maintain cellular senescence, the p53-p21-pRB and the p16-pRB pathways (Sultana et al., 2018). Induction of either of these converging pathways results in cell cycle arrest. Along with cell cycle arrest, senescent cells show a host of key phenotypes. These include resistance to apoptosis, reduced lysosomal function, increased senescence-associated β-galactosidase (SA-βGal) expression, lipofuscin accumulation, increased nuclear/cytoplasm ratio, loss of laminin B1, HMGB1 re-localization, secretion of damage- associated molecular pattern (DAMPs), damaged mitochondria, increased ROS formation, oxidative protein damage, and the SASP [Reviewed in (Martinez-Zamudio et al., 2017)]. SASP is the secretion of inflammatory cytokines, chemokines, growth factors, and proteases. The activation of SASP causes a local inflammation and, if the senescent cell is not removed by the immune system, is harmful to surrounding cells (Acosta et al., 2013).

Special AT-Rich Sequence-Binding Protein 1 (SATB1) is a transcriptional regulator which has recently been identified as a risk factor for PD (Chang et al., 2017; Nalls et al., 2018). Previously, we have reported that the activity of SATB1 is reduced in the vulnerable brain region of PD patients (Brichta et al., 2015). SATB1 is a ubiquitously expressed chromatin organizing factor that confers cell type-specific patterns of transcription and mediates a transcriptional response of midbrain DA neurons to toxic insult.

Here we show that the genetic ablation of SATB1 induces a senescence phenotype in hESC derived DA neurons. In DA neurons lacking SATB1, we observed the hallmarks of cellular senescence. We found that SATB1 directly represses the expression of p21, a crucial determinant of cellular senescence. Inhibition of p21 in SATB1 knockout (KO) cells ameliorates the senescence phenotype. Strikingly, elimination of SATB1 from cortical neurons did not induce senescence, or p21 expression. *In vivo*, the reduction of SATB1 in dopamine neurons caused p21 elevation and a local immune response. Finally, we found that p21 is actively expressed in the dopamine neurons of the SNpc of sporadic PD patients, rendering these cells prone to enter a state of senescence.

## Results

### hESC Derived DA Neurons Require SATB1

We first compared the role of SATB1 in the PD vulnerable neuronal population, DA neurons, to a less PD vulnerable neuronal cell type, CTX neurons (Giguere et al., 2018; Pedersen et al., 2005; Rajput and Rozdilsky, 1976). To produce pure DA neuronal cultures, we used an optimized DA neuron differentiation protocol (Figure 1A) (adapted from (Kriks et al., 2011; Miller et al., 2013; Steinbeck et al., 2015)), with the goal of obtaining high-purity DA neuron cultures. To achieve this high purity, we generated a NURR1 (a postmitotic marker for DA neurons (Saucedo-Cardenas et al., 1998))-driven GFP expressing hESC line (NURR1-GFP cells). Using a CRISPR/Cas9 based knock-in approach we replaced the endogenous NURR1-stop codon with P2A-H2B-GFP (Figure S1A). These NURR1- GFP cell derived DA neurons co-express GFP with both NURR1 and tyrosine hydroxylase (TH) antibody staining (Figure S1B). We then used FACS-based isolation of NURR1+ cells at day 25 of differentiation (Figure S1D). This enabled us to isolate a pure population of GFP+ cells with excellent survival (Figure S1C-D). Single cell qRT-PCR data in differentiated NURR1-GFP cells demonstrate that upregulation of DA neuron markers occurs between day 25 and day 40 of differentiation. These data also confirmed that 97% of the resulting NURR1-GFP cell-derived DA neurons were TH+ (Figure 1A, S1E). These data show that the sorted NURR1+ cells are highly pure and have the molecular characteristics of DA neurons (Figure 1A). To generate a SATB1 knockout in these cells, we used CRISPR/Cas9 to genetically eliminate SATB1 from human embryonic stem cells (hESCs) and differentiated them into the relevant neuronal subtypes. Sanger sequencing confirmed the successful mutagenesis (Figure S1F). SATB1 protein was eliminated from the hESC SATB1^KO^ cell line, which retained normal cellular appearance in culture (Figure 1B,C).

Using this approach, we differentiated SATB1^KO^ NURR1-GFP cells into mature DA neurons. Following floor plate differentiation, SATB1^KO^ NURR1-GFP cells expressed key markers such as OTX2, FOXA2 and LMX1A. These markers define the cells as midbrain floor plate precursors at day 16 of differentiation (Figure 1D). To track the functional maturation of the hESC-derived DA neurons, we performed electrophysiological recordings at different time points. Neither WT nor SATB1^KO^ neurons showed spontaneous action potentials at day 30 of differentiation. Cells of both genotypes on day 36 of differentiation started to display spontaneous action potentials, evoked action potentials, as well as voltage-depended Na+ and K+ currents (Figure S1G). Finally, the frequency of action potentials was equally increased at day 40 of differentiation (Figure 1E). These results show that the differentiated KO cells exhibited features of mature DA neurons. However, SATB1^KO^ neurons were unable to maintain the response to positive current injections (Figure 1F).

The knockdown of SATB1 from the midbrain was shown to be sufficient to cause degeneration of DA neurons in mice (Brichta et al., 2015). We therefore monitored the survival of SATB1^KO^ DA neurons during maturation. SATB1^KO^ DA neurons showed a significant reduction in survival starting at day 40 of differentiation (Figure 1G). Interestingly, we observed a plateau in cell viability which was maintained over the 60 days of differentiation. Surviving SATB1^KO^ DA neurons showed significantly decreased neurite outgrowth and complexity at day 60, following normal development at earlier days of differentiation (Figure 1H).

To determine the cell type-specificity of this finding, we differentiated the same KO hESC cells into CTX neurons. After this differentiation protocol, both WT and SATB1^KO^ expressed the relevant differentiation markers of layer V CTX neurons (Figure 1I). During the course of differentiation, survival of SATB1^KO^ CTX neurons was unaffected, and both neurite outgrowth and complexity showed a slight delay in development, but both were still able to achieve wild-type levels (Figure J-K). To verify SATB1 expression in our CTX neuronal cultures, we compared SATB1 protein level between DA and CTX neurons. We found that SATB1 is still abundant in CTX neurons, though to a lesser extent than in DA neurons (Figure 1L). Taken together, these findings suggest that the loss of SATB1 has separate physiological effects in these neuronal subtypes.

### SATB1 Has Discrete Regulatory Roles in DA and CTX Neurons

To understand the distinct functions of SATB1 in DA and CTX neurons, we performed concurrent RNA-Seq and ChIP-Seq experiments (Figure 2A). We used ChIP-Seq to compare the genome-wide binding profiles of SATB1 in early DA neurons, mature DA neurons and CTX neurons. We found that SATB1-binding had the highest intensity in mature DA neurons. This is reflective of the higher protein level (Figure 1L) and indicates an important role of the protein in this cell type (Figure 2B). We confirmed this finding by analysis of the expression profile changes caused by SATB1^KO^ in DA and CTX neurons.

**Figure 1.**
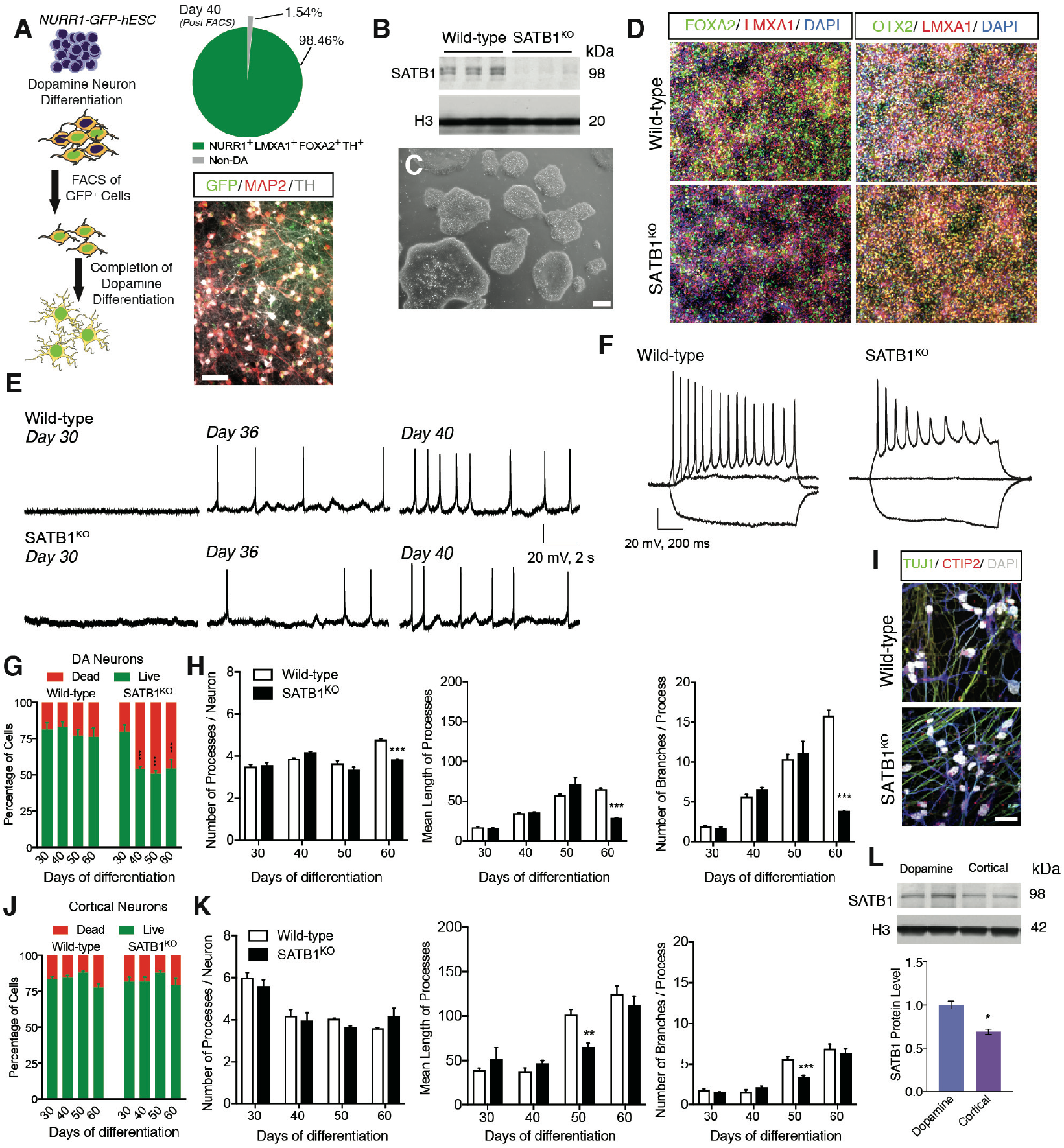
hESC Derived DA Neurons Require SATB1 to Maintain their Identity. (A) Overview of the differentiation protocol to generate highly enriched cultures of mature DA neurons. Distribution of DA neuron population post FACS, as determined by single-cell qRT-PCR. Immunofluorescent staining of the sorted NURR1-GFP+ neurons with anti-TH and anti-MAP2 showing the high yield of TH+ neurons following differentiation. (B) Western blot analysis to confirm SATB1^KO^ in hESC on protein level. (C) Microscopic bright light image of SATB1^KO^ hESC shows normal appearance. Scale bar: 200 μm. (D) Immunolabeling of DA precursor markers in WT and SATB1^KO^ hESC derived DA neurons after 16 days of differentiation reveals no difference in marker expression. (E) Depiction of spontaneous action potentials of WT and SATB1^KO^ DA neurons at different time points show that both genotypes differentiate into mature DA neurons between day 35 and 40. Scale bar: 20 mV, 2 s. (F) DA neurons at day 40 show significant differences in maintenance of response to positive current injections. Scale bar: 20 mV, 200 ms. (G) Longitudinal comparison of cell survival of WT vs. SATB1^KO^ DA neurons revealed a significant reduction in SATB1^KO^ survival between day 30 and 40, reaching a plateau at ~50%. Data are represented as mean ± SEM. (H) Quantification of neurite morphology and complexity in WT and SATB1^KO^ DA neurons. Data are represented as mean ± SEM. (I) Triple immunolabeling of cortical shows that both WT as well as SATB1^KO^ neurons express the essential markers for cortical neurons. Scale bar: 50 μm. (J) SATB1^KO^ cortical neurons show no significant change in survival during differentiation. Data are represented as mean ± SEM. (K) Quantification of neurite morphology and complexity in WT and SATB1^KO^ CTX neurons. (L) Representative western blot and quantification depicts that WT DA neurons express significantly higher levels of SATB1 than WT cortical neurons, suggesting a higher demand in DA neurons. Data are represented as mean ± SEM. See also Figure S1.

Comparison of WT and SATB1^KO^ DA neurons at an early timepoint (day 30) revealed few changes in gene expression (Figure 2C). At this timepoint, the cells were phenotypically comparable to WT. At day 50 of differentiation, when surviving SATB1^KO^ neurons show a phenotype, much greater gene expression changes were observed (Figure 2D). CTX neurons showed very few expression changes caused by SATB1^KO^ (Figure 2E). Principal components analysis of expression data can distinguish the DA neurons based on both their maturation and SATB1 genotype. The KO of SATB1 has a more dramatic effect in mature DA neurons than in early DA neurons. The analysis could not distinguish CTX neurons by SATB1 genotype (Figure S2A). We then evaluated the overlap in genes altered by SATB1^KO^. We found that there was almost no overlap in gene expression changes between the DA and CTX neurons (Figure S2B), nor shared GO pathway enrichment (Table S1). Next, we used the binding and expression target analysis (BETA) software (Wang et al., 2013) to incorporate the ChIP-Seq and RNA-Seq data. This analysis showed that SATB1 has no significant effects as gene regulator in early DA neurons (Figure 2F). In mature DA neurons SATB1 acts as gene repressor (p= 0.000236) (Figure 2G). In CTX neurons, it has both repressive (p=0.00791) and activating (p=2.97e-05) functions (Figure 2H), although SATB1 regulates far fewer genes in this cell type (Figure 2E).

**Figure 2.**
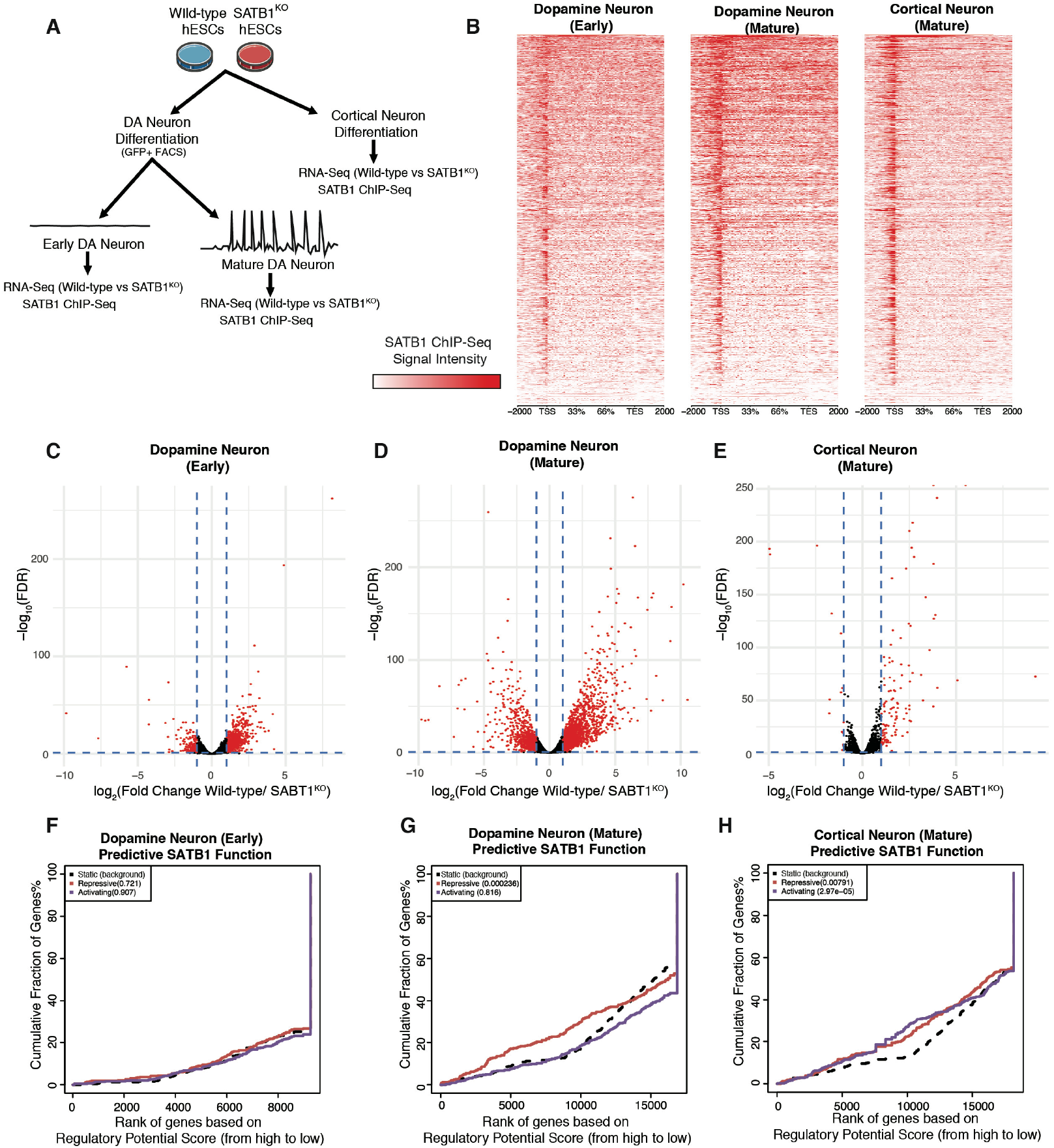
SATB1 Plays Discrete Regulatory Roles in DA and CTX Neurons. (A) Outline of the experimental approach comparing expression and DNA-binding and regulator profile of SATB1 in DA and CTX neurons. (B) Genome-wide heatmaps of SATB1-ChIP-Seq experiments comparing binding patterns in early and mature DA and CTX neurons. RNA-Seq expression profile comparing WT vs. SATB1^KO^ of early DA neurons (C), mature DA neurons (D), and mature cortical neurons (E). Red dots indicate significantly changed genes (FDR < 0.05, > ±2-fold expression change) BETA plots of combined computational analysis of SATB1-ChIP-Seq and RNA-Seq data of early DA neurons (F), mature DA neurons (G), and mature cortical neurons (H). Black line: static background, red line: repressive function, blue line: activating function. See also Figure S2.

### Loss of SATB1 in Dopamine Neurons Results in a Senescence Phenotype

Amongst the enriched GO pathways of SATB^KO^ DA neurons, we found an enrichment of the cellular senescence pathway. This pathway was not enriched in cortical knockout cells. This DA neuron enrichment was confirmed by GSEA of the mature SATB1^KO^ DA neuron transcriptome (Figure 3A). Given this, we sought to investigate if SATB1^KO^ DA neurons present the classical features of cellular senescence. First, we observed a dramatic increase in acidic lysosomal senescence associated beta- Galactosidase (SA-βGal) activity. This is the hallmark senescence biomarker (Figure 3B). Another key feature of senescent cells is the activation of the SASP. To determine if SATB1^KO^ DA neurons present this phenotype, we evaluated the expression of the described key SASP factors (Coppe et al., 2008). We found an upregulation of the majority of the SASP factors after 50 days of differentiation in the SATB1^KO^ DA neurons compared to WT neurons (Figure 3C). We confirmed SASP activation by western blotting. In the conditioned media of SATB1^KO^ neurons we found IGFBP7, which was absent in the media of WT neurons (Figure 3D). In fact, secretion of IGFBP7 alone is capable of inducing cellular senescence of surrounding cells (Severino et al., 2013). The expression data did not reveal any SASP activation in CTX SATB1^KO^ neurons (Figure S3). Another well described phenotype of cellular senescence is an increase in nucleus diameter correlating with reduced laminin B1 expression. Using automated high-content imaging, we evaluated nucleus size of SATB1^KO^ DA neurons in comparison to controls. We found a consistent, and significant increase in nucleus diameter in the DA neurons lacking SATB1 (Fig. 3G). We also found a corresponding significant decrease in lamin B1 expression (Figure 3E). Remarkably, CTX SATB1^KO^ neurons show neither of these nuclear features (Figure S3). Senescence-associated mitochondrial dysfunction leads to the elevation of reactive oxygen species. This in turn leads to oxidative protein damage. Using an OxyBlot assay, we found that SATB1^KO^ DA neurons show increased oxidized proteins compared to WT (Figure 3E). Furthermore, we found electron dense lipofuscin accumulations in the cytoplasm of SATB1^KO^ DA neurons (Figure 3F).

**Figure 3.**
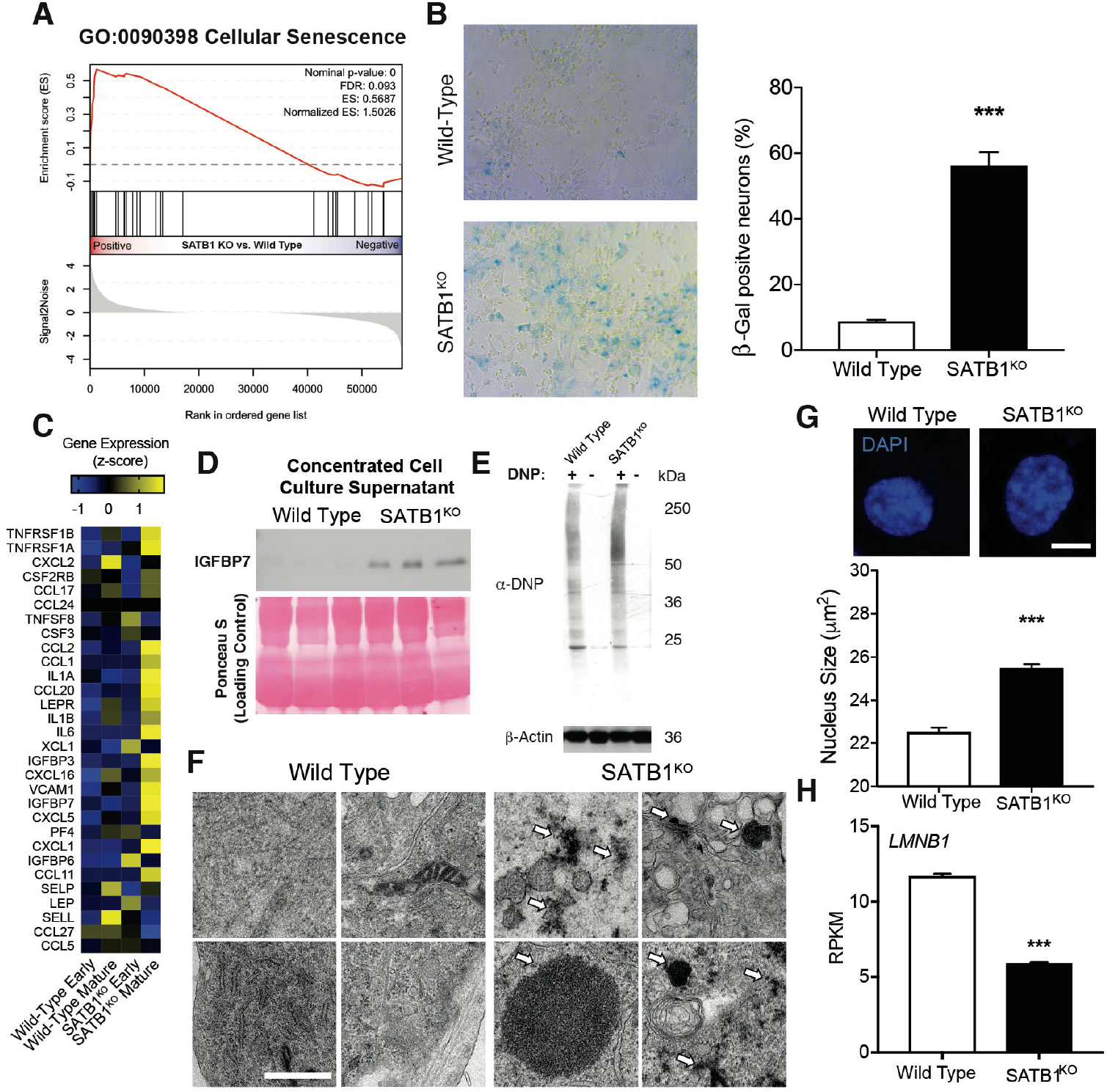
Loss of SATB1 in Dopamine Neurons Results in a Neuronal Senescence Phenotype. (A) GSEA plot of gene enrichment significantly correlating with the expression profile of cellular senescence. (B) Representative microscopic overview images of WT and SATB1^KO^ neurons subjected to X-Gal staining. Blue cells are senescence cells. Bar graph shows significant increase of SA-βGal positive cells. Data are represented as mean ± SEM. (C) Heatmap of genes associated with SASP. (D) Western blot of cell culture supernatant of mature DA neurons. The SASP factor IGFBP7 is secreted from SATB1^KO^ neurons. Ponceau S stain serves as loading control. (E) Oyxblot-assay shows increased protein oxidation in SATB1^KO^ neurons. (F) Representative TEM-image of the electron-dense lipofuscin observed in SATB1^KO^ neurons. (G) Representative images from high-quantity imaging of nuclei. (H) RNA-Seq expression of LMNB1 in SATB1 DA neurons. Scale bar: 5 μm Bar graph shows significant increase of nucleus size in SATB1^KO^ neurons. Data are represented as mean ± SEM. See also Figure S3.

### Dopamine Neurons Lacking SATB1 have impaired Lysosomal and Mitochondrial Function

SA-βGal activity is indicative of altered function and enlargement of lysosomal compartments. Analysis of RNA-Seq data show a drastic change in lysosomal gene expression in SATB1^KO^ DA neurons (Figure 4A). Based upon these molecular alterations and because of the reported connections between lysosomal alterations and senescence we performed a detailed characterization of lysosomal function.

**Figure 4.**
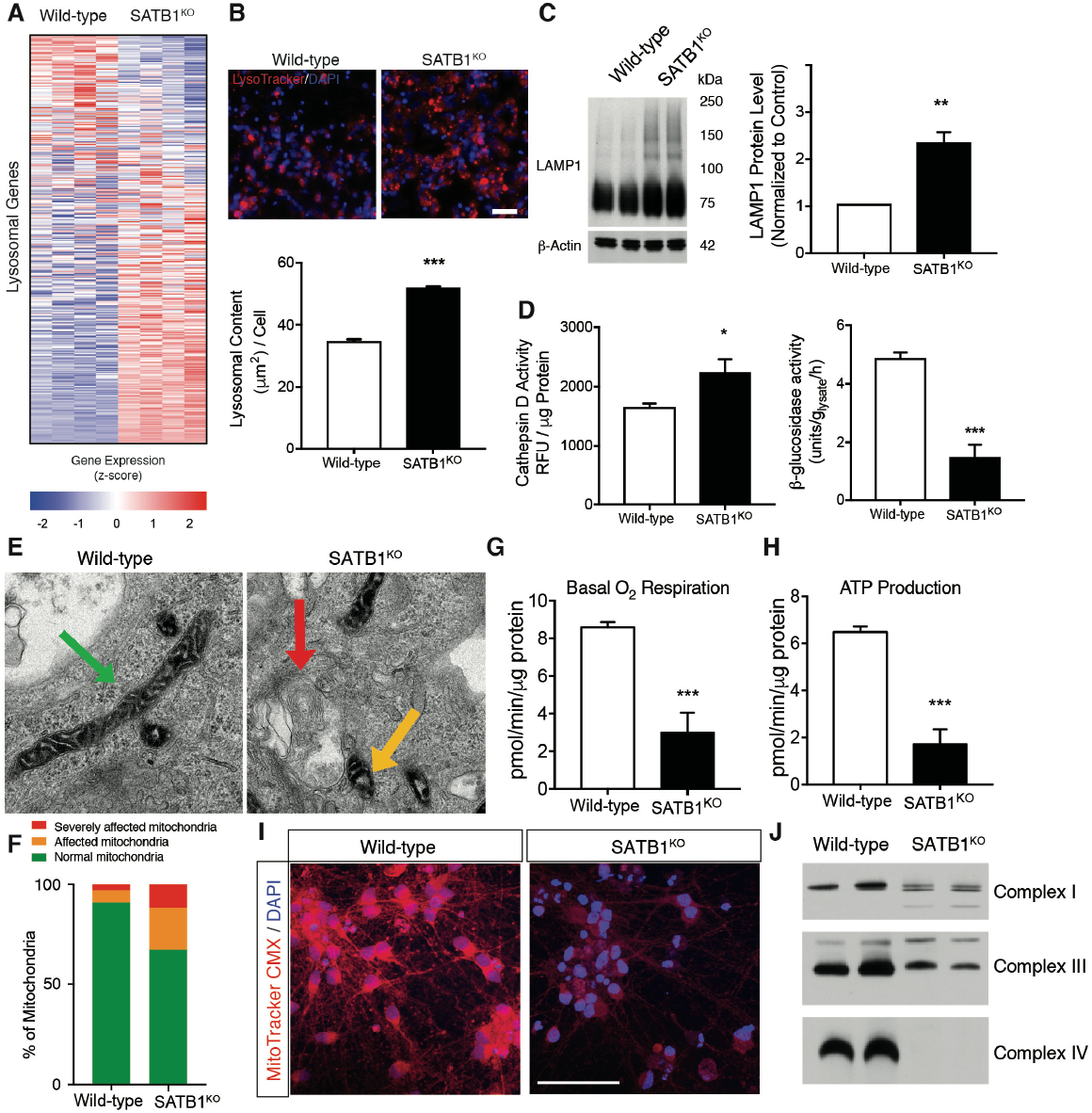
Dopamine Neurons Lacking SATB1 have impaired Lysosomal and Mitochondrial function. (A) Heatmap of the RNA expression of lysosomal genes comparing WT and SATB1KO neurons. (B) Representative confocal images of LysoTracker staining of WT and SATB1^KO^ neurons. Bar graph shows significant increase of lysosomal content in SATB1^KO^ neurons. Scale bar: 100 μm. Data are represented as mean ± SEM. (C) Representative western blot of mature DA neurons. Bar graph shows significantly increased LAMP1 levels in SATB1^KO^ neurons. Data are represented as mean ± SEM. (D) Bar graphs of cathepsin D and GCase assays. Data are represented as mean ± SEM. (E) Representative TEM-images of mitochondria in mature DA neurons. Green arrows mark normal, yellow marks affected and red arrow marks severely affected mitochondria, quantified in (F). Analysis of cellular respiration identified reduced basal respiration (G) and ATP production (H) in SATB1KO neurons. Data are represented as mean ± SEM. (I) Representative images of MitoTracker CMX stain shows decreased membrane potential in SATB1^KO^ neurons. (J) Native-PAGE of mitochondrial complex integrity using anti-NDUFA9 (Complex I), anti-UQRC2 (Complex III) and anti-MTCO2 (Complex IV) antibodies. See also Figure S4.

Using LysoTracker dye and automated high-content imaging, we found that SATB1^KO^ DA neurons showed a significant increase in endo/lysosomal content, a well characterized feature of senescent cells (Figure 4B). Importantly, the endo/lysosomal content of SATB1^KO^ CTX neurons remained unchanged (Figure S3A). To confirm the increase in endo/lysosomal content, we quantified the protein level of the endo/lysosomal marker LAMP1. In line with the imaging data, LAMP1 levels were significantly increased in SATB1^KO^ DA neurons (Figure 4C). Given the structural, ultrastructural (Figure S4) and molecular changes in the lysosomes of SATB1^KO^ DA neurons, we evaluated the lysosomal function in these cells. To assess lysosomal degradation capacity, we used two representative enzymatic activity assays which play important roles in lysosomal degradation. Both the cathepsin D and the β-glucosidase activity assays were significantly altered, implicating severe lysosomal dysfunction (Figure 4D).

Since the demonstration of an increase in oxidized proteins suggested mitochondrial dysfunction, we assessed the function of these organelles in SATB1^KO^ DA neurons. We first quantified ultrastructural alterations in the mitochondria of SATB1^KO^ DA neurons by electron microscopy (EM). Doing so, we found that these neurons have more damaged and abnormal mitochondria (Figure 4E, F). The evaluation of functioning mitochondria using MitoTrackerCMX, and oxygen consumption, by Seahorse respirometry, revealed a significant reduction in basal respiration and ATP production in SATB1^KO^ DA neurons (Figure 4G, I). Importantly, cellular respiration and ATP production remained unchanged in SATB1^KO^ CTX neurons (Figure S3B,C). In addition, KO of SATB1 in CTX neurons did not alter components of the mitochondrial complexes. Moreover, the assembly of mitochondrial complexes was found to be greatly impaired in SATB1^KO^ DA neurons. This was also reflected in the total protein level of the complexes which were altered in SATB1^KO^ DA neurons, although to a lesser extent (Figure 4J, S4).

### SATB1 Repression of CDKN1A Prevents Senescence in DA Neurons

We next sought to determine a direct link between SATB1 and the observed cellular senescence phenotype in SATB1^KO^ DA neurons. To this end, we checked SATB1 DNA binding in the regulatory regions of critical factors regulating cellular senescence. Using ChIP-Seq in DA neurons, we found that SATB1 binds the regulatory region of *CDKN1A* (Figure 5G). This gene encodes for p21, a critical regulator of senescence. Both *CDKN1A* transcription and p21 protein levels were significantly increased in SATB1^KO^ DA neurons (Figure 5A-C), together signifying a repressive function of SATB1 at this locus. The elimination of SATB1, and the resulting de-repression of *CDKN1A*, is thus sufficient to elevate cellular p21 levels. In line with these findings, SATB1 has previously been described to regulate cellular senescence in keratinocytes (Lena et al., 2012), and has also been previously described to repress the promoter of *CDKN1A* (Wang et al., 2014).

**Figure 5.**
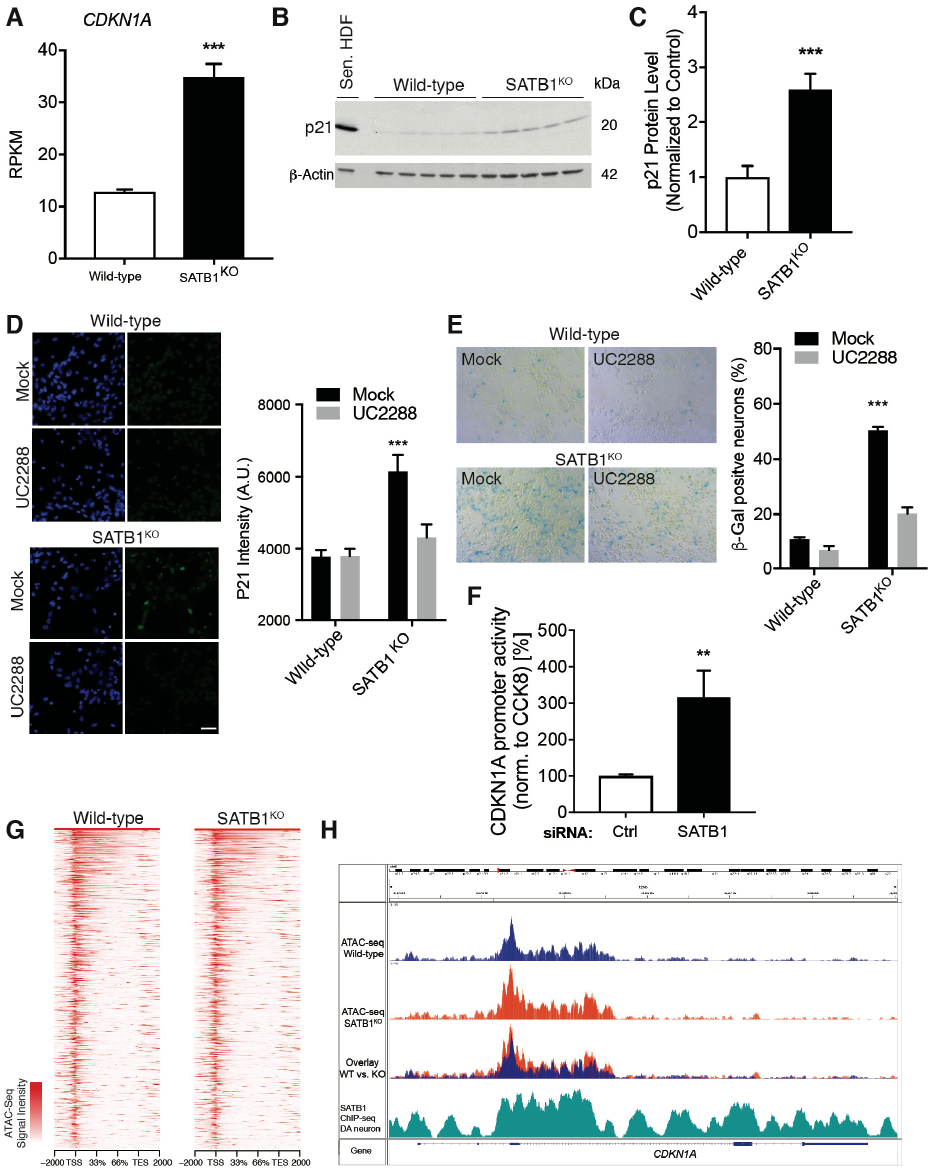
SATB1 Repression of CDKN1A Prevents Senescence in DA Neurons. (A) Bar graph shows significant upregulation of *CDKN1A* expression. Data are represented as mean ± SEM. (B) Western blot probed for p21 and b-actin comparing WT and SATB1^KO^ DA neurons. Sen. HDF: senescence induced fibroblasts as positive control. (C) Bar graph shows significant upregulation of p21 level. Data are represented as mean ± SEM. (D) Representative confocal images of p21 staining of WT and SATB1^KO^ neurons with and without UC2288 treatment. Scale bar: 10 μm. Bar graph shows significant increase of p21 level in SATB1^KO^ neurons which is normalized by UC2288 treatment. Data are represented as mean ± SEM. (E) Representative microscopic images of WT and SATB1^KO^ neurons subjected to X-Gal staining with and without UC2288 treatment. Bar graph shows significant increase of percentage of SA-βGal positive SATB1^KO^ neurons which is normalized by UC2288 treatment. Data are represented as mean ± SEM. (F) Luciferase reporter assay for the *CDKN1A* promoter under conditions of SATB1 knockdown in HeLa cells. (G) Heat maps of ATAC-Seq experiments comparing open chromatin patterns in WT and SATB1^KO^ mature DA neurons. (H) The *CDKN1A* gene overlaid with ATAC-Seq enrichment tracks from WT and SATB1^KO^ DA neurons as well as SATB1-ChIP-Seq from WT DA neurons. See also Figure S5.

We next sought to determine the direct role of p21 in inducing the senescent phenotype in SATB1^KO^ DA neurons. To do so, we used the small molecule UC2288 which decreases cellular levels of p21 (Wettersten et al., 2013). Indeed, UC2288 treatment for five days significantly reduced p21 levels in SATB1^KO^ DA neurons (Figure 5D). p21 elimination in SATB1^KO^ DA neurons, by UC2288 treatment, was able to significantly reduce the number of SA-βGal positive cells (Figure 5E). This suggests that elevated p21 mediates the senescent phenotype in SATB1^KO^ DA neurons. To determine if SATB1 directly represses *CDKN1A*, we performed a luciferase reporter assay, and found that under conditions of SATB1 knockdown in HeLa cells, *CDKN1A* promoter activity was significantly increased (Figure 5F). To further confirm the specificity of the *CDKN1A* regulation by SATB1, we performed ATAC-Seq experiments in WT and SATB1^KO^ DA neurons. The ATAC-Seq approach revealed that the global open chromatin structure is not changed in SATB1^KO^ cells (Figure 5G). This indicates a specific mode of gene regulation by the protein in these cells. Indeed, comparing the *CDKN1A* locus between WT and SATB1^KO^ DA neurons, the ATAC-Seq shows a more opened gene region which overlaps with the SATB1 ChIP-Seq binding profile (Figure 5G).

Interestingly, SATB1^KO^ causes an increase of p21 protein level in DA neurons but did not change *CDKN1A* transcription or protein levels of p21 in CTX neurons (Figure S5 A-B). Importantly, we found that the *CDKN1A* locus is more open in WT DA neurons compared to CTX neurons, suggesting a DA neuron-specific role for this gene (Figure S5C). This finding implies an additional mechanism of transcriptional regulation of p21. During the process of p21-mediated cellular senescence, transcription factors such as SMAD3 or FOXO1 increase *CDKN1A* gene expression (Martinez-Zamudio et al., 2017). Both SMAD3 and FOXO1 play crucial regulatory roles in DA neurons (Doan et al., 2016; Tapia-Gonzalez et al., 2011). Correspondingly, their expression is higher in DA compared to CTX neurons (Figure S5D, E).

We hypothesize that *CDKN1A* gene expression in DA neurons is under constant influence of promoting factors which aim to increase cellular p21 levels, whereas in CTX neurons this is not the case. SATB1 is therefore an important repressor of *CDKN1A* transcription (Wang et al., 2014) and regulates p21 levels in DA neurons.

### In vivo Reduction of SATB1 Induces Senescence of DA Neurons

Next, in order to validate the human *in vitro* findings, we shifted to an *in vivo* mouse model. In this model, we sought to investigate whether the reduction of SATB1 triggers p21 expression and subsequent senescence in vivo. Using stereotactic adeno-associated virus 1 injection expressing short hairpin RNA (AAV1-shRNA), we downregulated the endogenous SATB1 in the midbrain of mice (Figure 6A). As previously reported, this knockdown of SATB1 results in an elimination of TH+ neurons three to four weeks after injection (Brichta et al., 2015). We investigated the midbrain of these mice two weeks after injection to investigate expression changes and found only a reduction of TH+ DA neurons (Figure S6A). Analyzing DA neuron-specific bacTRAP expression data (Brichta et al., 2015), we found in Satb1- knockdown mice the second highest upregulated gene *Cdkn1a* and an increase in genes related to the SASP (Figure 6C,E). Moreover, when we stained the remaining DA neurons for expression of p21, we found a significant increase in p21 positive DA neurons (Figure 6D). To assess whether SASP in DA neurons triggers an immune response in the midbrain, we stained for the microglial marker Iba-1. Strikingly, confocal microscope analysis of Satb1-knockdown midbrains two weeks after virus injection revealed that microglia are co-localizing with the TH+ DA neurons, suggesting a release of immune factors to attract immune cells (Figure S6B). This mechanism is in line with the finding of significantly elevated expression levels of immune factors (Figure 3C, 6E). To confirm that the SASP in these animals activated the surrounding microglia, we adapted a previously reported method to characterize the activation status of microglia based upon their roundness and cell size (Davis et al., 2017). Using this approach, we found a significantly increased percentage of activated microglia in Satb1-knockdown substantia nigra (Figure 6F), which target Satb1 depleted DA neurons (Figure S6B,C).

**Figure 6.**
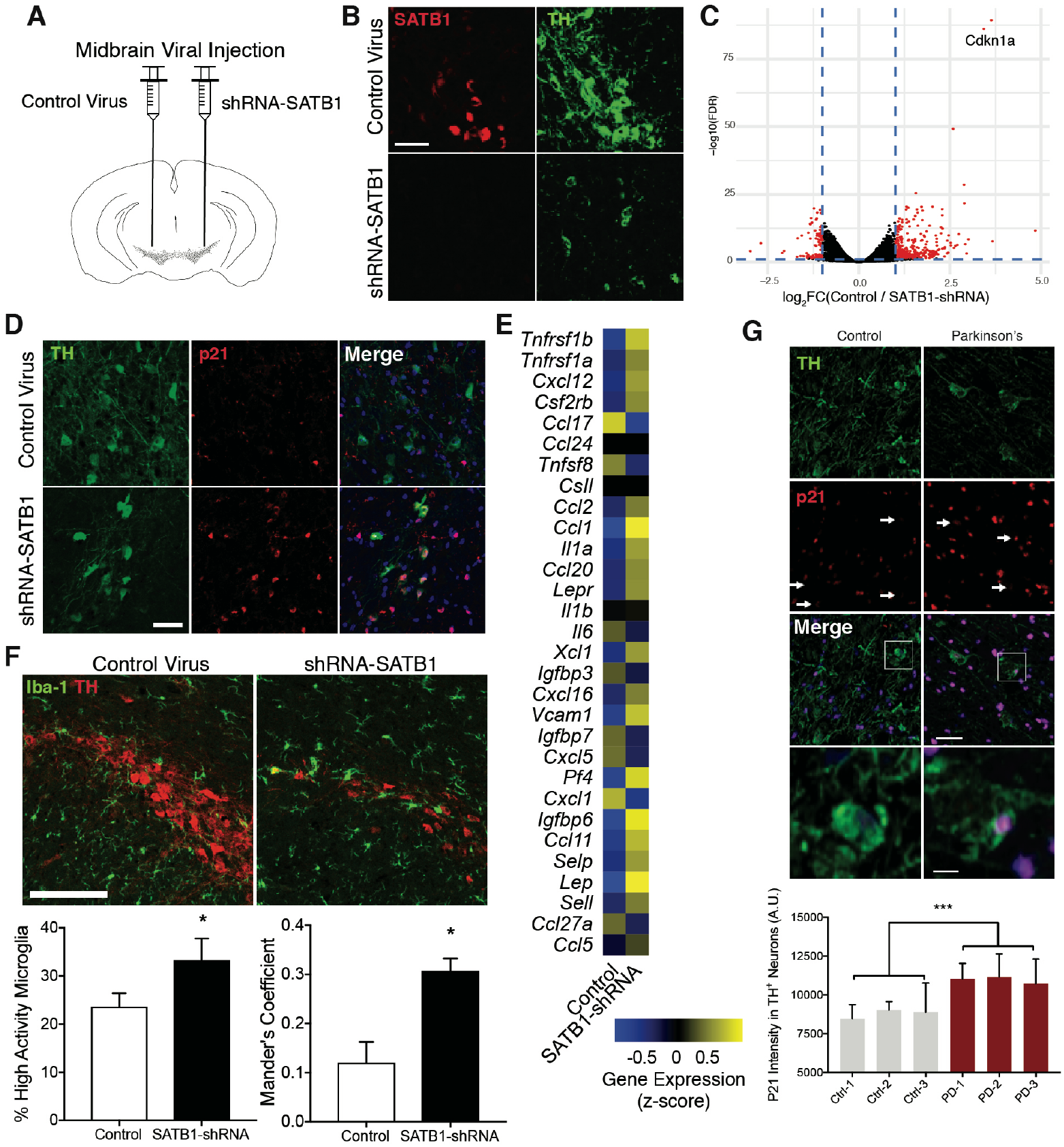
In vivo reduction of SATB1 induces senescence of DA Neurons. (A) Virus injection strategy. (B) Representative confocal images of TH and Satb1 staining of control-shRNA and Satb1-shRNA TH+ neurons in the SNpc, 2 weeks after injection. Scale bar: 50 μm (C) DAT-bacTRAP-based expression profile changes in DA neurons comparing control-shRNA and Satb1-shRNA. Cdkn1a amongst the highest upregulated transcripts. (D) Representative confocal images of TH and p21 staining of control- shRNA and Satb1-shRNA TH+ neurons in the SNpc, 2 weeks after injection. Scale bar: 50 μm (E) Heatmap of mouse genes associated with SASP. (F) Representative confocal images of TH and Iba-1 staining of control-shRNA and Satb1-shRNA TH+ neurons in the SNpc, 2 weeks after injection. Scale bar: 100 μm Quantification of high activity microglia, and colocalization between Iba-1 and TH signals, in the SNpc, comparing control-shRNA and Satb1-shRNA. Data are represented as mean ± SEM. (G) Representative confocal images of TH and p21 staining of human brain slices of the SNpc derived from age-matched control individual or sporadic PD patient. (H) Quantification of intensity of p21 signal in TH+ SNpc neurons of three age-matched control individuals and three PD patients Scale bar: 50 μm, 5 μm. Data are represented as mean ± SEM. See also Figure S6.

Using a regulatory network analysis of expression data from substantia nigra samples of human subjects with incipient PD, we have previously reported that the regulatory function but not the expression level of SATB1 is significantly decreased (Brichta et al., 2015; Zheng et al., 2010). Based on this finding and the above described discoveries, we analyzed the expression of p21 in human sporadic PD patient midbrain slices (obtained from the Harvard Brain Tissue Resource Center). Crucially, we found a significant increase in the intensity of p21 staining within TH+ DA neurons in PD patient brain slices compared to age-matched controls (Figure 6G,H). This finding potentially links aberrant SATB1 regulation of *CDKN1A* to PD.

## Discussion

The general understanding of cellular senescence is tied to its initial discovery in mitotic cells more than five decades ago by Hayflick and Moorhead (1961). Hayflick and Moorhead discovered that primary cells in culture undergo a limited number of cell divisions. These cells then reach a state of replicative senescence and can no longer regenerate tissue (Hayflick, 1965). Since then, cellular senescence and the associated SASP has been thoroughly investigated and many additional biological roles have been identified. Senescence has an established function in cancer prevention by triggering growth arrest and by signaling to the immune system to remove incipient cancerous cells (Acosta et al., 2013; Acosta et al., 2008; Georgilis et al., 2018; Kang et al., 2011). In addition, senescence and SASP have been shown to contribute to development and tissue repair (Munoz-Espin and Serrano, 2014). If immune cells do not remove senescent cells, their constant pro-inflammatory SASP can be toxic to surrounding cells and therefore has been associated with multiple age-related diseases (Franceschi and Campisi, 2014). Recent studies have suggested that not only mitotic, but also postmitotic cells are capable of entering a state of senescence. This has been observed in a number of cell types from the CNS (Jurk et al., 2012), bone (Farr et al., 2017), cardiac (Wang et al., 2016), and adipose tissue (Minamino et al., 2009).

Evidence suggests that neurons can enter a state of cellular senescence (Baker and Petersen, 2018; Tan et al., 2014). A number of pathological features in the brains of Alzheimer‘s and Parkinson’s disease patients overlap with phenotypes observed in cellular senescence (Tan et al., 2014). Whether neurons contribute to these features is an open question. Indeed, some studies revealed progressive activity of SA-βGal in cultured cerebellar granular neurons and hippocampal neurons (Bhanu et al., 2010; Bigagli et al., 2016; Green et al., 2017). However, SA-βGal activity by itself has been debated to be a sufficient marker for cellular senescence (Severino et al., 2000).

In the present study, we have utilized pure human stem cell-derived DA neurons, model mice and sporadic PD patient brain slices to establish that SATB1 is a regulator of cellular senescence in DA neurons. We show that SATB1^KO^ DA neurons show virtually all established features of cellular senescence. This includes activation of the SASP, SA-βGal activity, lysosomal dysfunction, lipofuscin accumulation, damaged and dysfunctional mitochondria, enlargement of the nucleus, increased oxidative protein damage, and expression pathway changes towards cellular senescence. Interestingly, we did not observe increased DNA damage in the senescent cells. However, we found that SATB1 binds directly to the regulatory region of *CDKN1A* repressing its expression, which is in accordance with previous findings (Wang et al., 2014). Moreover, inhibition of *CDKN1A* in SATB1^KO^ DA neurons significantly reduced the number of senescent cells, demonstrating that the direct regulation of p21 levels in SATB1^KO^ DA neurons triggers the senescence phenotype. Strikingly, when the SATB1^KO^ stem cell clone was differentiated into CTX neurons, none of the senescence phenotypes were observed.

The SATB1-dependent repression of *CDKN1A* transcription seems crucial to DA neuron function. Loss of SATB1 in DA neurons caused a ~300% elevation in CDKN1A transcription, but did not significantly alter levels in CTX neurons. It is tempting to speculate that the CDKN1A locus is under a constant transcriptional activation pressure in DA neurons compared to CTX neurons, which is reflected by significantly higher expression of *CDKN1A*-promoting factors (Figure S5 D,E; Tinkum et al., 2013). These findings suggest that the intrinsic properties of DA neurons, be they structural (size, branching), energetic demand, or high stress level (Pacelli et al., 2015) may render these cells vulnerable to cellular senescence.

As previously reported, the transcriptional activity of SATB1 decreases in incipient PD patient brains (Brichta et al., 2015). This suggests that a p21-induced senescence-like phenotype may be present in human PD brain. Indeed, we describe here a significant increase in p21 levels in DA neurons in the SNpc of PD patients in comparison to age-matched controls. There are a number of studies which revealed phenotypes in PD brains which could be attributed to cellular senescence (Tan et al., 2014): the discovery of elevated inflammatory factors in the CSF of PD patients for example could be linked to SASP (Blum-Degen et al., 1995). Other studies found evidence for dysfunctional lysosomes in the PD brain (Chu et al., 2009; van Dijk et al., 2013). Also, the accumulation of lipofuscin, increased SA-βGal activity, and enlargement of mitochondria has been described in PD (Braak et al., 2003; Trimmer et al., 2000; van Dijk et al., 2013). However, none of the above-mentioned studies have connected these findings to neuronal senescence (Tan et al., 2014). Evidence of a p21-dependent cellular senescence in PD is supported by genetic mouse models of the disease. Neural stem cells from Prkn−/− mice show an increase of p21 protein levels (Park et al., 2017). In line with this finding, LRRK2 mutant animals show robust increases in *Cdkn1a* transcript levels (Nikonova et al., 2012). These findings further link familial forms of PD to the p21 pathway.

PD is an age-related disorder. Even familial cases of PD will usually develop the disease later in life. Age is therefore considered the most important contributing risk factor for PD. Importantly, the accumulation of senescent cells is also clearly linked to increased age (Soto-Gamez and Demaria, 2017). In young individuals, senescent cells which show a SASP will be detected and eventually removed by the immune system. Yet, with age, as immune surveillance decreases, senescent cells can escape the immune system, a phenomenon directly linked to aging (van Deursen, 2014). Prolonged presence of senescent cells in the tissue is problematic since the SASP secrets pro-inflammatory factors which can trigger local inflammation and spread the senescence phenotype in a paracrine fashion (Acosta et al., 2013). Should cellular senescence be a step preceding the loss of SNpc cells during the development of PD, this would explain why disease symptoms occur before the loss of soma in the midbrain. DA neurons which enter senescence would lose function but would remain in the midbrain until microglia remove them. This would be similar to what we observed in our Satb1-knockdown mouse model (Figure 6F). In aged PD patients, it is possible to envision that senescent DA neurons would not be removed. Instead, they would cause a local inflammation and spreading of senescence, resulting in the expression of the respective markers, which indeed have been identified in PD brains (Tan et al., 2014).

The ultimate confirmation that cellular senescence contributes to aging and age-related disorders came from animal studies which showed that removal of senescent cells has a beneficial effect on the disease phenotype and lifespan (Baker et al., 2016; Baker et al., 2011; Xu et al., 2018). These discoveries are driving the field of the development of drugs which reduce aging by removal of senescent cells. These drugs, termed senolytics, have already been able to improve age-related phenotypes in mice (Xu et al., 2018; Zhu et al., 2015). Based on our finding that SATB1 regulates cellular senescence in DA neurons, we hypothesize that SATB1 itself could be a promising target for novel senolytics. Moreover, therapeutic strategies targeting SATB1 or p21 in PD may be a beneficial route to intervention.

## Materials and Methods

For description of materials and methods, see supplemental materials.

## Author Contributions and Notes

M.R., B.K., T.W.K., J.C., J.N., J.A.P, E.J.P., K.D., D.A., G.C., W.W. and J.Z. performed experiments and/ or analyzed data. M.R., B.K., T.W.K, D.A., L.S.R-E, L.S., and P.G. planned the project and supervised experiments. J.S. provided materials for the project. M.R., B.K., T.W.K., L.B., L.S., and P.G. wrote the manuscript.

## Acknowledgments

We thank the Genomics Resource Center and the Electron Microscopy Resource Center (EMRC) at Rockefeller University for technical support. All mitochondrial respiration measurements were performed using the Agilent Seahorse XFe96 analyzer, as well as High-Content Imaging on the ImageXpress system with generous support from the Rockefeller University High-Throughput Screening and Spectroscopy Resource Center (HTSRC). Human brain tissue was provided by the Harvard Brain Tissue Resource Center, which is supported in part by PHS grant number R24 MH068855. This work was supported by the United States Army Medical Research and Material Command (USAMRMC) under Award No. W81XWH-10-1-0640 (LB), W81XWH-12-1-0039 (MR)and the JPB Foundation, Award #475 (PG). T.W.K. is supported by a NYSCF Druckenmiller fellowship. Opinions, interpretations, conclusions and recommendations are those of the authors and are not necessarily endorsed by the sponsors.

